# Altered Duodenal Mucosa-Associated Microbiota and Immune Profiles in Functional Dyspepsia: A Study of Host-Microbiome Homeostasis

**DOI:** 10.1101/2025.04.10.648308

**Authors:** Emily C. Hoedt, Grace L. Burns, Seungha Kang, Jessica Bruce, Mark Morrison, Simon Keely, Nicholas J. Talley

## Abstract

Recent work suggests an altered duodenal mucosa-associated microbiota (MAM) in patients with functional dyspepsia (FD) when compared to outpatient controls, these differences may reflect alterations in host-microbiome homeostasis. Given that specific mucosal immune signatures have been identified in FD, we hypothesised that these changes would be associated with specific microbial changes. We aimed to profile the duodenal MAM to identify microbes associated with known changes in FD mucosal and peripheral immune homeostasis. Duodenal biopsies were collected from 11 asymptomatic outpatient controls and 17 FD patients. Separate biopsies were collected for MAM 16S rRNA amplicon sequencing, histological evaluation, and mucosal lamina propria mononuclear cell (LPMC) isolation. Where available peripheral blood mononuclear cells (PBMC) were isolated from whole blood. PBMC and LPMC populations were analysed for CD4 and CD8 T cell populations by flow cytometry. Initial comparisons of histological and immune measures revealed significant differences between FD and outpatient controls, with decreased villi goblet cells and increased LPMC CD4 Central Memory, LPMC CD8, and PBMC CD4+ Central Memory Th17 in FD compared to controls. While microbiome profiles varied between groups, specific associations with the histological and immunological scoring found that in controls, villi goblet cells correlated positively with *Massilia* and negatively with *Exiguobacterium*, while FD patients showed no significant correlations. Additionally, controls exhibited a negative correlation between LPMC CD4 Central Memory and *Veillonella*, with FD patients showing no significant correlation. Notably, FD patients demonstrated a significant negative correlation between LPMC CD8 and *Sulfophobococcus*, and a positive correlation between PBMC CD4+ Central Memory Th17 and both *Gemella* and *Fusobacterium*.

**Importance:** While several papers have reported the alteration in FD duodenal MAM, study numbers are low and consensus on specific biomarker signatures within the microbiome have not been reached. Our findings contribute to this growing body of evidence, indicating that patients with FD exhibit distinct alterations in duodenal MAM and immune profiles compared to outpatient controls. Furthermore, the immune-microbiome associations present in control populations were absent in FD patients suggesting a loss of host-microbiome homeostasis that may contribute to FD pathophysiology. Our work highlights potential microbial biomarkers which may be a consequence or driver of the mucosal micro-inflammation state which is a characteristic of FD. Future work is required to validate whether these specific microbes are responsible for driving inappropriate host immune responses to inform treatment strategies and assist with disorder diagnosis.

## Introduction

Functional dyspepsia (FD) is a highly prevalent disorder of gut-brain interaction (DGBI), affecting 6.8% of the global population, based on Rome IV criteria (1). Individuals suffering from FD experience postprandial fullness, early satiation, epigastric pain, and epigastric burning (2). While no overt structural or biochemical signatures identifiable though routine clinical workup have emerged, a number of subclinical physiological and immune signatures have been identified, suggesting a loss of mucosal homeostasis in individuals with FD (3, 4). A subset of patients with FD have been found to have high duodenal eosinophil counts (5–11), and increased proportions of effector T cells including Th2 and Th17-like cells in the duodenum and decreased duodenal goblet cell numbers (12). These features have been referred to as “microinflammation” in the literature (11) and suggests that it is antigen-driven, potentially driven by the microbiome *per se* and/or exposure to specific food antigens.

In that context, the duodenum is recognised to harbor mucosa-associated microbiota (MAM) and with specific signatures evident in both asymptomatic and FD patients (13–18) and with several microbial taxa associated with FD symptoms. For instance, increases in the relative abundance of Fusobacteria (14, 18) and genera such as *Alloprevotella*, *Corynebacterium*, *Peptostreptococcus*, *Staphylococcus*, *Clostridium*, and *Streptococcus* (16, 18, 19) have been reported. Conversely, decreased relative abundance of genera *Actinomyces*, *Gemella*, *Haemophilus*, *Megasphaera*, *Mogibacterium*, and *Selenomonas* has been observed (14, 16, 18–20). Given that the duodenum is one of the initial sites where food antigens are exposed to the immune system, these microbial shifts may contribute to the dysregulation of immune homeostasis in FD. However, it is unclear if this is a cause or consequence of disease and whether there are functional implications associated with this MAM signature.

There are a number of ways in which disturbances to microbiota homeostasis could influence the mucosal immune system. The loss of key homeostasis-promoting commensals may facilitate the outgrowth of potential pathobionts, or as is suspected in the case of *Faecalibacterium prausnitzii* in IBD (21, 22), the loss of immune-regulatory microbes may have a permissive effect on the immune system, resulting in inappropriate responses. Further, alterations in the microbiome may lead to differential digestion of food antigens (23), driving aberrant immune responses. Here, we hypothesised that specific immune features in FD are linked to changes in the MAM and aimed to identify microbes associated with peripheral and mucosal immune signatures across the oesophagus, stomach and duodenum of FD patients. We found that key associations between duodenal immune cells and specific microbial genera that were present in outpatient controls were lost in patients with FD. Our findings further support the concept that loss of host-microbiome homeostasis is a feature of FD pathology and may contribute to patient symptoms.

## Methods

### Study recruitment

Participants were recruited and sampled as previously described (3). In summary participants aged 18-80 years were recruited through outpatient gastroenterology clinic at John Hunter Hospital in Newcastle, New South Wales, Australia. Patients met the Rome III criteria for postprandial distress syndrome (PDS), or epigastric pain syndrome (EPS) with or without PDS (EPS ± PDS). Given that post-prandial epigastric pain largely contributes to overlap between the Rome III subtypes (24, 25), and we had small numbers of ‘pure’ EPS subjects, we pooled these groups to create the EPS ± PDS subgroup, as previously published (12). Asymptomatic outpatient controls were patients requiring endoscopy for routine care, such as for unexplained iron deficiency anaemia (IDA), positive faecal occult blood test (+FOBT), gastroesophageal reflux disease (GERD) or dysphagia, with no organic GI disease confirmed during endoscopy. Exclusion criteria for the study included patients with a body mass index (BMI)>40, organic GI conditions and pregnant women.

### Ethics statement

The studies involving human participants were reviewed and approved by Hunter New England (reference 13/12/11/3.01). The patients/participants provided their written informed consent to participate in this study.

### Sample collection

As previously described (3), at endoscopy upper gastrointestinal biopsies were collected using standard biopsy forceps from 11 asymptomatic outpatient subjects and 17 FD patients. Oesophageal, stomach, and duodenal biopsies were stored in RNAlater at −20ᵒC until processing.

### DNA extraction

Samples were first lysed through bead homogenization (3mins at 5,0000rpm) using the Precellys24 after removal from RNAlater solution as previously described (26). The total DNA was then extracted and purified with the Promega Maxwell automated DNA recovery system following manufacturer’s instructions.

### 16S rRNA amplicon gene sequencing

The 16S rRNA gene encompassing the V6 and V8 regions was targeted using the 917F (5’-GAATTGRCGGGGRCC;-3’) and 1392wR (5’-ACGGGCGGTGWGTRC;-3’) primers were modified to contain Illumina specific adapter sequence as previously described (26). The 16S library was constructed following the Illumina protocol #15044223 Rev.B (Illumina, Inc., United States) by the Australian Centre for Ecogenomics (ACE). Indexed amplicons were pooled together in equimolar concentrations and sequenced on MiSeq Sequencing System (Illumina) using paired end sequencing with V3 300bp chemistry. Passing quality control (QC) of resulting sequence was determined as 10,000 raw reads per sample prior to data processing and passing QC metrics in line with Illumina supplied reagent metrics of overall Q30 for 600bp reads of >70%.

### Microbiome analysis

A total of 81 samples were sequenced under one sequence run. Data was processed with QIIME2 (v2020.11) and DADA2, as previously described (27, 28). Taxonomic assignment was performed in R (v4.1.3) with DADA2 (v1.20.0) against SILVA SSU r138 reference database. Analysis of microbiota diversity was performed in R, using packages phyloseq (v1.38.019), breakaway (v4.7.3), ampvis2 (v2.7.620), and microbiome (v1.16.0). Rare taxa that had less than 0.01 percent relative abundance across all samples were removed. Alpha diversity was analysed for significant difference through Wilcoxon test, Bonferroni correction was used for post-hoc comparisons. Adonis2 (PERMANOVA) was used to test statistical significance of Bray-Curtis PCoA. Data was normalised with total sum scaling (TSS) and subsequently transformed using centre log ratio (CLR) for rank sum testing.

### Predictive Functional Profiling

Functional predictions of 16S rRNA gene amplicons were completed with PICRUSt2 using QIIME2 plug-in (v2021.11) and annotated with the MetaCyc database (https://metacyc.org/). Data was normalised and transformed as described above using R and visualized using packages phyloseq, SIAMCAT (v1.12.0), microbiome, and microbiomeutilities (v1.00.16).

### Phenotypic immune and histological data

Histology, peripheral blood mononuclear and lamina propria mucosal cells (PBMC and LPMC, respectively) data were obtained from previously published work (3).

### Data availability

16S rRNA amplicon raw sequence data are available and have been deposited under NCBI BioProject accession number PRJNA1229776. NCBI SRA record is accessible with the following link http://www.ncbi.nlm.nih.gov/bioproject/1229776.

## Results

### Patient demographics

Biopsies for MAM sequencing were available for 11 asymptomatic outpatient controls (mean age 58; 36% female; 18% FOBT, 27% IDA, 45% dysphagia, and 18% reflux) and 17 FD patients (mean age 46; 72% female). There was some variation in biopsy availability across the upper GI sites: oesophagus (controls n=10; FD n=16), stomach (controls n=11; FD n=17), and duodenum (controls n=10; FD n=17). Additionally, PBMC samples were available for 6 out of 11 controls and 11 out of 17 FD patients, while duodenum LPMC samples were available for 8 out of 10 controls and 14 out of 17 FD patients.

### Comparison of histological and immune characteristics suggests immune alterations

For those patients which had samples available for MAM analyses we next compared the immune status in FD and controls for duodenum histology and flow evaluated cells with them LPMC and PBMC populations (**Table 2**). We observed a significant decrease in FD villi goblet cells (P=0.05) compared to controls. Additionally, there was a significant increase in CD4 Central Memory cells (P=0.01) and CD8 cells (P=0.04) in the LPMC population of FD patients. In PBMC, FD patients showed a significant increase in CD4+ Central Memory Th17 cells (P=0.02). These findings suggest distinct immune alterations in FD patients, particularly in memory T cell populations.

**Table 1.** Demographics for functional dyspepsia (FD) and control cohort with available samples for microbiome analyses. Body mass index (BMI); proton pump inhibitors (PPI); epigastric pain syndrome (EPS); postprandial distress syndrome (PDS); iron deficiency anaemia (IDA), positive faecal occult blood test (FOBT), gastroesophageal reflux disease (GERD).

|  | Control (n= 11) | FD (n= 17) | P-value |
| --- | --- | --- | --- |
| Sex (M/F) | 7\4 | 4\13 | 0.053 |
| Age (median) | 61 | 47 | 0.07 |
| BMI | 26 | 26 | 0.63 |
| PPI | 2 (2 Unknown) | 8 (4 Unknown) | 0.18 |
| **FD subtypes** |  |  |  |
| EPS |  | 2 |  |
| PDS |  | 7 |  |
| EPS/PDS |  | 8 |  |
| **non-FD subtype** |  |  |  |
| IDA | 4 |  |  |
| FOBT | 2 |  |  |
| GERD | 2 |  |  |
| Dysphagia | 3 |  |  |
| IBS diagnosis | 0 | 4 | 0.13 |
| *H. pylori* positive | 1 | 0 | 0.39 |
| **Eosinophil status** |  |  |  |
| low | 9 | 13 | 0.99 |
| high | 2 | 4 | 0.99 |
| **Smoking history** |  |  |  |
| Current | 2 | 3 | 0.99 |
| Ex | 4 | 7 | 0.99 |
| Never | 3 | 6 | 0.99 |

**Table 2.** Comparison of immune status in FD vs controls for 1) histology evaluated duodenal cells 2) flow cytometry evaluated cells. High-power fields (hpf); lamina propria mononuclear cell (LPMC); peripheral blood mononuclear cells (PBMC); * denotes significant result when groups were compared with unpaired t-test.

| Histology- duodenum | Control (n=11) | Patient (n=17) | P-value |
| --- | --- | --- | --- |
| Eosinophils (per hpf) | 18.91±8.78 | 17.24±8.88 | 0.38 |
| IELS (per 50 enterocytes) | 11.82±6.24 | 16.94±21.64 | 0.65 |
| Degranulation score | 1.91±0.83 | 1.82±1.02 | 0.60 |
| Gland goblet cell | 13.45±5.50 | 9.59±6.10 | 0.12 |
| Villi goblet cells | **10.73±4.45** | **6.88±4.23** | **0.05\*** |
| CD117 Mast cells (per hpf) | 9.72±2.42 | 9.70±1.79 | 0.97 |
| Total goblet cells | 25±10.21 | 15.94±10.13 | 0.06 |
| LPMC- duodenum | Control (n=8) | Patient (n=14) | P-value |
| CD3 | 37.90±13.38 | 41.66±16.04 | 0.71 |
| CD4 | 31.71±10.87 | 36.29±13.50 | 0.44 |
| CD4 Naive | 1.34±1.04 | 3.46±5.98 | 0.28 |
| CD4 Effector | 3.58±2.81 | 5.081±3.71 | 0.48 |
| CD4 Effector Th1 | 0.19±0.53 | 1.23±2.41 | 0.26 |
| CD4 Effector Th2 | 23.51±21.22 | 19.67±14.01 | 0.93 |
| CD4 Effector Th17 | 37.05±24.43 | 48.12±22.96 | 0.30 |
| CD4 Central Memory | **1.50±0.99** | **4.22±4.21** | **0.01\*** |
| CD4 Central Memory Th1 | 4.83±5.24 | 7.14±8.744 | 0.71 |
| CD4 Central Memory Th2 | 33.20±18.92 | 40.24±18.57 | 0.40 |
| CD4 Central Memory Th17 | 14.29±9.57 | 16.67±9.54 | 0.27 |
| CD4 Effector Memory | 1.004±0.96 | 1.78±1.85 | 0.33 |
| CD4 Effector Memory Th1 | 7.26±6.73 | 9.57±10.04 | 0.63 |
| CD4 Effector Memory Th2 | 14.82±10.62 | 20.64±13.71 | 0.22 |
| CD4 Effector Memory Th17 | 16.38±14.80 | 15.53±11.22 | 0.99 |
| CD8 | **12.23±7.20** | **19.74±7.09** | **0.04\*** |
| CD8 Naive | 0.97±0.74 | 2.09±3.06 | 0.27 |
| CD8 Effector | 0.68±0.48 | 0.57±0.24 | 0.62 |
| CD8 Central Memory | 2.07±1.94 | 3.98±2.35 | 0.06 |
| CD8 Effector Memory | 0.33±0.31 | 0.70±0.62 | 0.09 |
| PBMC | Control (n=6) | Patient (n=11) | P-value |
| CD3+ | 41.97±24.09 | 31.13±20.82 | 0.35 |
| CD4+ | 38.21±19.83 | 37.95±17.93 | 0.53 |
| CD4+ Naive | 9.20±11.94 | 12.44±15.18 | 0.57 |
| CD4+ gut homing | 0.63±1.04 | 0.85±1.56 | 0.56 |
| CD4+ Effector | 1.66±3.08 | 1.91±2.47 | 0.21 |
| CD4+ Effector Th1 | 1.17±2.87 | 3.50±5.13 | 0.22 |
| CD4+ Effector Th2 | 19.17±20.13 | 32.08±29.05 | 0.36 |
| CD4+ Effector Th17 | 5.09±9.45 | 6.39±8.79 | 0.33 |
| CD4+ Central Memory | 9.70±15.33 | 3.33±4.06 | 0.98 |
| **CD4+ Central Memory Th1** | 6.71±9.34 | 11.31±10.01 | 0.40 |
| **CD4+ Central Memory Th2** | 18.98±17.90 | 13.12±13.25 | 0.61 |
| **CD4+ Central Memory Th17** | **4.20±5.65** | **12.95±10.68** | **0.02\*** |
| **CD4+ Effector Memory** | 0.95±1.47 | 0.98±1.56 | 0.42 |
| **CD4+ Effector Memory Th1** | 8.97±10.63 | 14.42±17.64 | 0.67 |
| **CD4+ Effector Memory Th2** | 2.93±5.11 | 1.85±3.29 | 0.84 |
| **CD4+ Effector Memory Th17** | 0.68±1.67 | 1.19±3.42 | >0.99 |
| **CD8+** | 8.80±15.16 | 17.59±14.80 | 0.09 |
| **CD8+ Naive** | 2.52±3.40 | 7.04±8.23 | 0.07 |
| **CD8+ gut homing** | 0.23±0.33 | 0.72±0.68 | 0.07 |
| **CD8+ Effector** | 2.64±6.16 | 2.82±3.43 | 0.14 |
| **CD8+ Central Memory** | 0.73±1.14 | 0.93±1.08 | 0.53 |
| **CD8+ Effector Memory** | 0.74±1.65 | 1.06±1.84 | 0.19 |

### FD patients have a decreased diversity predominately in the duodenum compared to outpatient controls

Comparisons between upper gastrointestinal sites microbiomes for outpatient “controls” and FD patients was first completed through alpha diversity indices Choa1, Shannon and Simpson at the ASV level (**Figure 1A)**. Although all measures indicated a decreased microbial diversity for FD patients compared to outpatient “controls” across all sample sites only the duodenum was significantly decreased for FD (Choa1 p-value 0.02, Shannon P=0.03 and Simpson P=0.04). In respect to beta diversity (PCoA Bray-Curtis) clustering and separation of microbial profiles was more evident for the stomach (PERMANOVA P=0.04) and duodenum (PERMANOVA P=0.02; **Figure 1B**).

**Figure 1.**
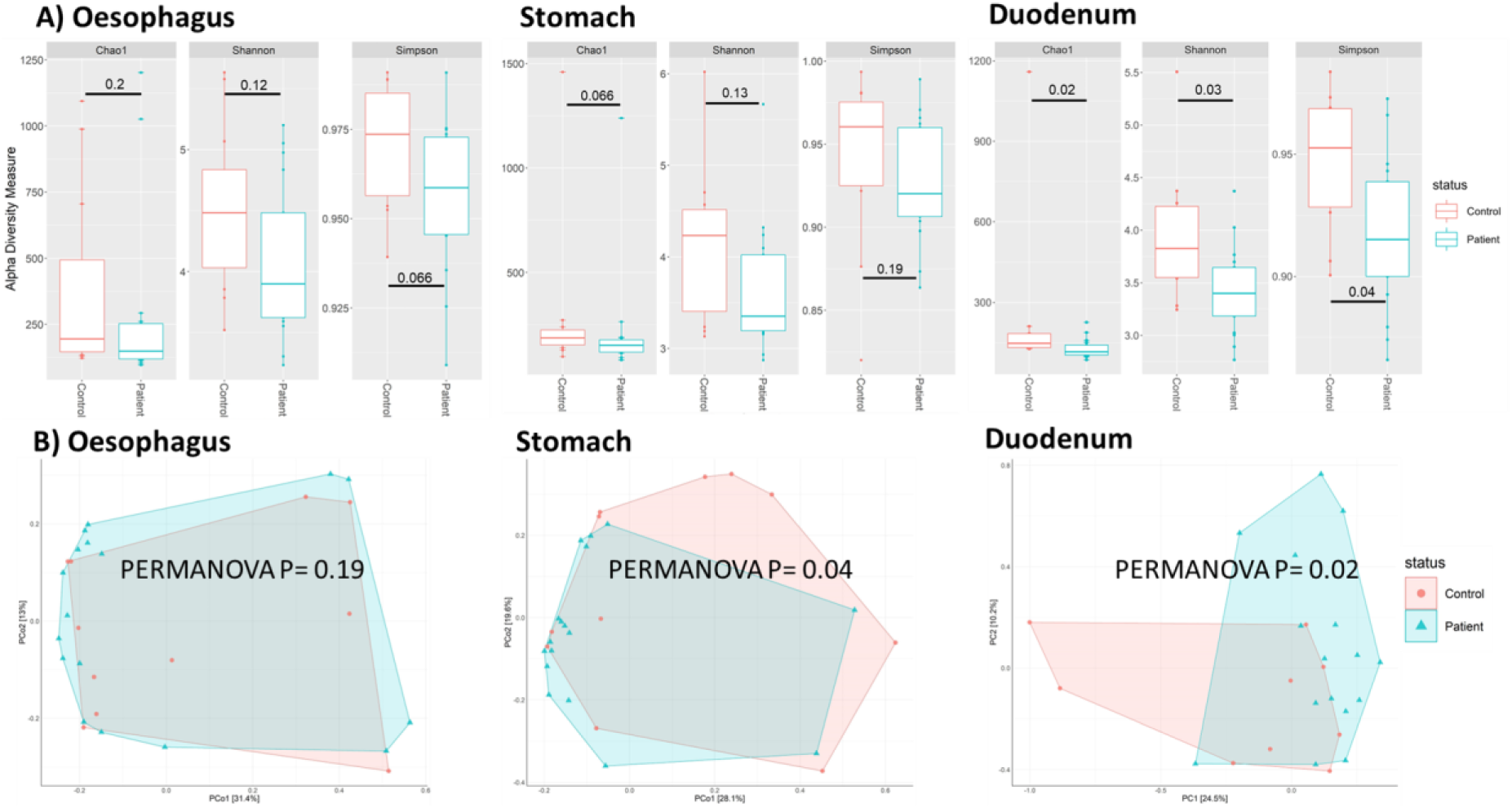
**A)** Alpha diversity indices Choa1, Shannon and Simpsons of controls (red) and functional dyspepsia patients (blue) for oesophagus, stomach, and duodenum. **B)** Beta diversity PCoA Bray-Curtis plot comparing controls (red) and functional dyspepsia patients (blue) for oesophagus, stomach, and duodenum respectively. Alpha diversity was analysed for significant difference through Wilcoxon test and beta diversity significance assessed by PERMANOVA P-values <0.05 were considered significant. Error bars represent standard deviation.

### Histology correlations – duodenum

Increased goblet cell activity can improve the presentation of antigens to T-cells, enhancing local immune surveillance and response (29, 30). However, FD patients have been reported to have a reduced number of duodenal goblet cell and mucin exocytosis (12), in addition to an increase in microinflammation with increased numbers of eosinophils reported (5–10). Therefore, we examined the correlations between duodenal MAM and previously reported duodenal eosinophil, goblet cell, and mast cell counts (3) (**Figure 2**). Only 2 genera were identified as being significantly associated with outpatient controls, *Massilia* was positively associated with gland, villi and total goblet cell counts, and *Exiguobacterium* was negatively associated with gland, villi and total goblet cell counts. Within FD patients only *Exiguobacterium* was positively associated with gland goblet cell counts.

**Figure 2.**
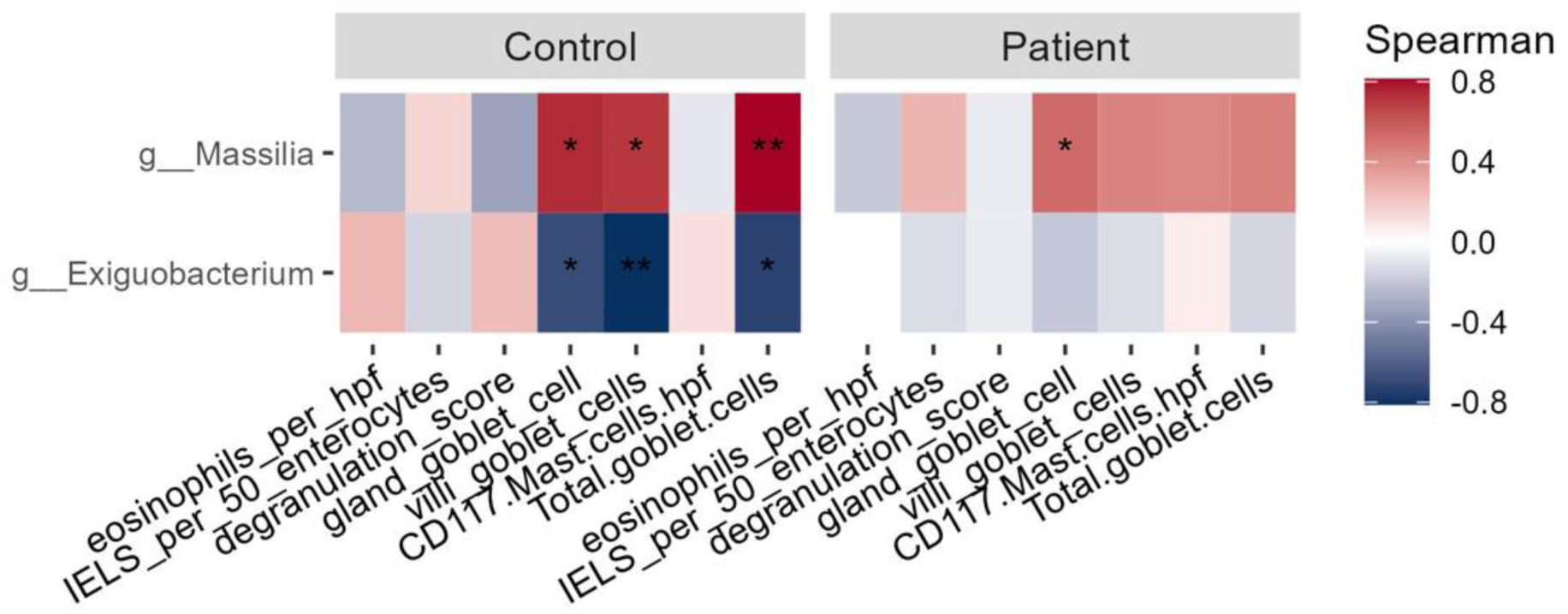
Spearman correlations for duodenal MAM genera and duodenal histological measures eosinophil per high-power fields (hpf), intraepithelial lymphocytes (IELs), degranulation score, gland goblet cell, villi goblet cell, total goblet cell counts and CD117 mast cell counts hpf. Red denotes a positive correlation and blue a negative correlation. Data was transformed with centre log ratio (clr). Adjusted P-values <0.05 were considered significant; *P<0.05, **P<0.01, ***P<0.001.

### Dysregulated effector and memory T cell response to duodenal MAM

Next, we examined the association between the duodenal MAM population and circulating peripheral effector and memory T cell profiles (CD3, CD4, and CD8) previously quantified within control and FD PBMCs (3). Differential correlative profiles were observed between groups, with 28 (22 positive) significant associations between genera and T cell profiles for outpatient controls, and 21 (8 positive) significant associations identified within FD patients (**Figure 3A**). Notably, multiple positive associations were observed across the CD4 effector and effector memory populations for *Oribacterium*, *Gemella*, and *Exiguobacterium* in outpatient controls. In contrast, the FD population exhibited a greater number of negative associations between genera and PBMC T cell profiles, particularly within the CD4 and CD8 populations. Unlike outpatient controls, FD patients showed positive associations with *Massilia* and *Cellulomonas* and CD3 T cell counts. CD3 T cells are responsible for activating both cytotoxic T cells (CD8+ naive T cells) and T helper cells (CD4+ naive T cells) (31).

**Figure 3.**
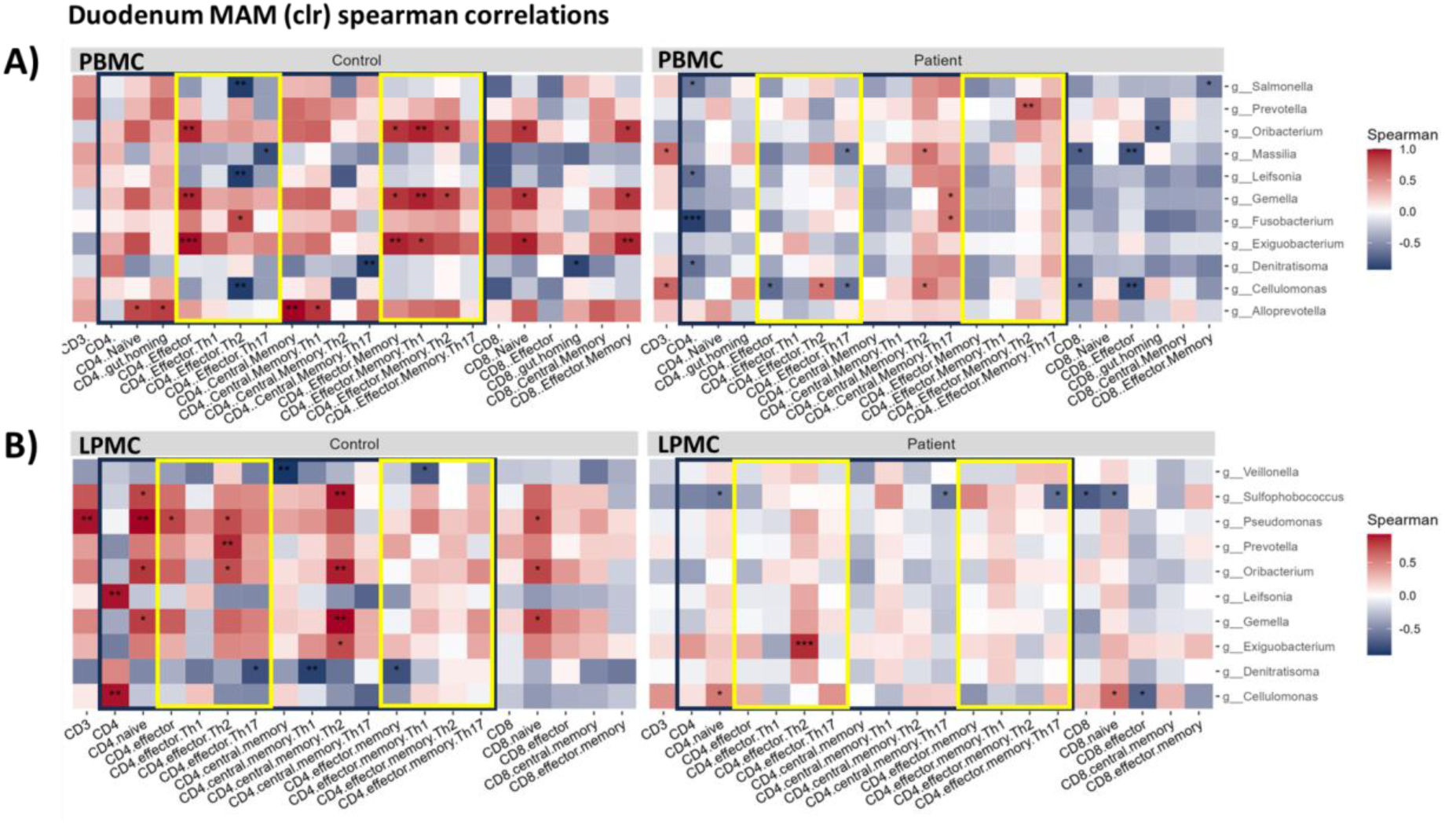
Spearman correlations for duodenum MAM genera, **A**) human peripheral blood mononuclear cells (PBMCs) and **B**) lamina propria mononuclear cells (LPMC) effector and memory T cell types quantified with flow cytometry. Black box groups all CD4 cell types and yellow boxes separate CD4 effector and effector memory cell types. Red denotes a positive correlation and blue a negative correlation. Data was transformed with centre log ratio (clr). Adjusted P-values <0.05 were considered significant; *P<0.05, **P<0.01, ***P<0.001.

Investigation of the lamina propria mucosal effector and memory T cells isolated from duodenal biopsies, and the duodenal MAM, revealed similarly disparate associative profiles between outpatient controls and FD patients (**Figure 3B**). Controls had a greater number of correlations (18/23 were positive correlations) compared to FD patients (3/9 positive correlations). Considering both the PBMC and LPMC effector and memory T cell profile associations with the duodenal MAM, these results suggest a dysregulated immune response in FD patients compared to outpatient controls in response to an altered duodenal MAM.

### MAM of different upper GI sites suggests altered peripheral effector and memory T cell response in FD patients

While no data was available for the lamina propria mucosal effector and memory T cells isolated from oesophagus or stomach biopsies we did analyse the correlation between the respective MAM upper GI sites and altered peripheral effector and memory T cell response. Similar to the duodenum (**Figure 3A**) the oesophagus FD correlative profile (24/48 positive significant correlations) appears to be disproportionate to the outpatient controls (36/44 positive significant associations; **Figure 4A**). Several genera in FD patients were negatively associated with CD3 *Granulicatella, Leptotrichia, Mogibacterium, Neisseria* and *Porphyromonas*, in addition to CD4 central memory cell types, *Desulfomicrobium, Exiguobacterium*, *Lentimicrobium*, *Lysobacter*, *Ottowia*, *Pseudomonas*, *Smithella*, *Staphylococcus*, TM7x and *Veillonella*. In contrast, outpatient controls had an abundance of different genera with positive correlations to CD4 effector, central memory and effector memory as well as CD8 cell types (**Figure 4A**).

**Figure 4.**
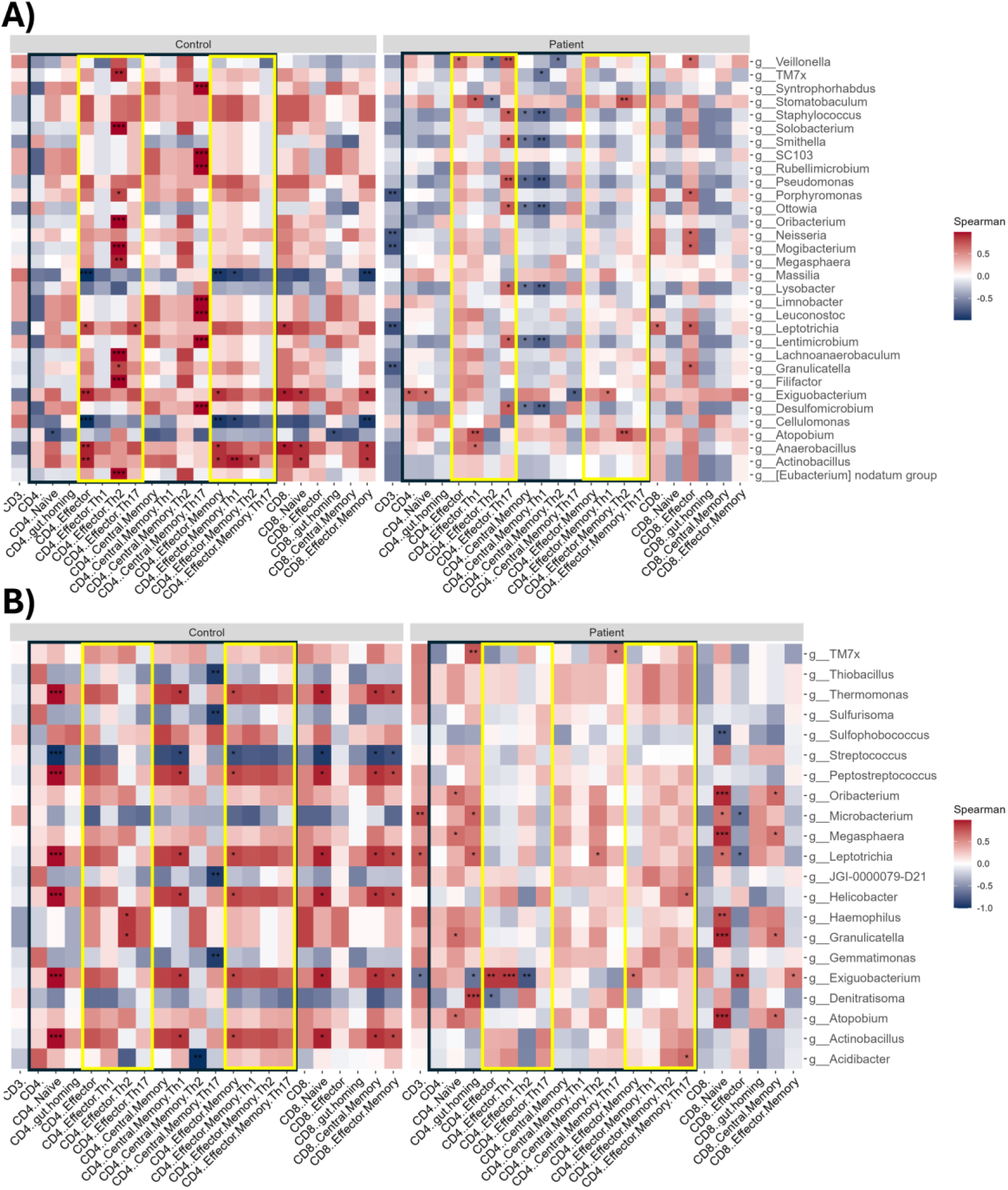
Spearman correlations for **A)** oesophagus **B)** stomach MAM genera, human peripheral blood mononuclear cells (PBMCs) flow cytometry markers. Black box groups all CD4 cell types and yellow boxes separate CD4 effector and effector memory cell types. Red denotes a positive correlation and blue a negative correlation. Data was transformed with centre log ratio (clr). Adjusted P-values <0.05 were considered significant; *P<0.05, **P<0.01, ***P<0.001.

Similarly, the correlations observed in the stomach suggest a differential response of gut homing T-cells and the MAM. We observed 38/49 positive correlations for our outpatient control cohort and 30/37 positive correlations for FD patients (**Figure 4B**). Seven genera were only associated with controls (positive correlations: *Actinobacillus*, *Peptostreptococcus*, *Thermomonas*; negative correlations: *Gemmatimonas*, JGI-0000079-D21, *Streptococcus*, and *Thiobacillus*) and nine with FD patients (positive correlations: *Acidibacter*, *Atopobium*, *Megasphaera*, *Oribacterium*, TM7x; negative correlation: *Sulfophobococcus*, and *Sulfurisoma*; mixed correlations: *Denitratisoma* and *Microbacterium*).

Across all PBMC MAM upper GI site correlations, only *Exiguobacterium* was consistently positively associated with peripheral effector and memory T cell types for outpatient controls, and six genera were shared across 2 MAM sites (positive: *Actinobacillus*, *Leptotrichia*, *Oribacterium*; negative: *Cellulomonas*, *Massilia*; mixed: JGI-0000079-D21). There was no conservation of genera associations across MAM sites and PBMC t cell types for FD patients, however there were six genera which were present across two upper GI sites (positive: *Atopobium*; mixed: *Denitratisoma*, *Exiguobacterium*, *Leptotrichia*, *Oribacterium*, and TM7x).

### Predicted MAM functional difference between FD and outpatient controls

Wilcox rank sum testing revealed significant differences in the predicted functions of the MAM across various gastrointestinal regions between outpatient controls and FD patients (**Table 3 and Table S1**). Specifically, four pathways were significantly different in the 133 pathways in the stomach, and 18 pathways in the duodenum. Two of the significantly different pathways from the oesophagus were also noted in the stomach. Furthermore, there were four significantly different pathways shared between the stomach and duodenum (**Table S1).** However, no significant Spearman’s correlations were found between these predicted functions and the histology or flow cytometry data.

**Table 3.** Top five (based on P-value) Wilcoxon rank sum testing results of significantly different PICRUSt2 predicted functional pathways between outpatient controls and FD patients across each upper gastrointestinal sampled mucosal associated microbiotas (MAMs) from centred log-ratio (CLR) transformed data. P-values calculated with Benjamini-Hochberg (BH) procedure, for controlling the False Discovery Rate (FDR).

| **Oesophagus MAM** | **MetaCyc pathway ID** | **BH FDR P-value** | **Increased in** |
| --- | --- | --- | --- |
| Norspermidine biosynthesis | PWY-6562 | 0.001 | Control |
| Mycothiol biosynthesis | PWY1G-0 | 0.036 | Control |
| P-cymene degradation | PWY-5266 | 0.047 | Patient |
| P-cumate degradation | PWY-5273 | 0.047 | Patient |
| **Stomach MAM** | **MetaCyc pathway ID** | **BH FDR P-value** | **Increased in** |
| Superpathway of sulfur oxidation (Acidianus ambivalens) | PWY-5304 | 0.001 | Patient |
| Pantothenate and coenzyme A biosynthesis I | PANTOSYN-PWY | 0.002 | Patient |
| L-histidine degradation I | HISDEG-PWY | 0.004 | Patient |
| Phosphopantothenate biosynthesis I | PANTO-PWY | 0.004 | Patient |
| P-cymene degradation | PWY-5266 | 0.004 | Patient |
| **Duodenum MAM** | **MetaCyc pathway ID** | **BH FDR P-value** | **Increased in** |
| Chondroitin sulfate degradation I (bacterial) | PWY-6572 | 0.011 | Patient |
| Adenosylcobalamin biosynthesis I (anaerobic) | PWY-5507 | 0.020 | Control |
| Superpathway of L-serine and glycine biosynthesis I | SER-GLYSYN-PWY | 0.020 | Patient |
| CMP-pseudaminic acid biosynthesis | PWY-6143 | 0.027 | Patient |
| Protein N-glycosylation (bacterial) | PWY-7031 | 0.027 | Patient |

## Discussion

Functional dyspepsia is considered a disorder of gut-brain interaction (DGBI), with symptoms commonly linked to psychological stress (32, 33). However, it is now clear that FD patients exhibit a range of sub-clinical physiological abnormalities in their duodenal mucosa, including increases in intestinal epithelial permeability, and mucosal immune cell numbers, as well as altered neuronal signalling and bile acid profiles (3–11, 34, 35). These changes coincide with consistent reports of an altered duodenal MAM, with FD patients exhibiting reduced microbiota diversity and changes in abundance of key genera (13–18). While it has been hypothesised that host-microbiome homeostasis is lost in FD, there is little direct evidence to show that these phenomena are related. Here we aimed to examine whether alterations in FD patient mucosal and peripheral immune profiles were associated with specific changes in the upper gastrointestinal tract MAM, including the oesophagus, stomach and duodenum. In our FD cohort, reduced alpha-diversity was evident in the duodenum, while beta-diversity was significantly altered in the stomach and duodenum. Importantly, FD patients had fewer associations between mucosal immune cell subsets and microbial genera than controls, coinciding with these differences in microbiota ecology. These data are the first to directly show a loss of host immune-microbiome homeostasis in patients with FD.

We have previously reported a reduction in duodenal goblet cells numbers in patients FD when compared to non-FD controls (12). The loss of goblet cells and resulting exocytosis likely impacts the overlying mucus-swelling microbes as changes in mucin secretion are known to impact the MAM (36). In contrast to controls, FD patients’ association between goblet cells and bacteria of the genus *Massilia* was not significant although still positively correlated (37, 38). Given the loss of mucus secreting cells in FD, the mucosal environment may be unfavourable to *Massilia* species. Another possibility, given that in FD patients the genus *Massilia* was positively associated with circulating Th2 memory cells, is that the altered mucosal environment in FD drives virulence factors in these microbes that provoke the immune system, such as reported opportunistic pathogens *Streptococcus pneumoniae* (39, 40) and *Escherichia coli* (41). Members of the genus *Massilia* are considered as opportunistic pathogens with increased *Massilia* abundance in the duodenum observed in Parkinson’s disease (42), pancreatic cancer (43), while infections of *Massilia* spp. have been found in immunocompromised patients (38, 44), in eye (37) and ear infections (45). In contrast to *Massilia*, the genus *Exiguobacterium* was negatively associated with goblet cells in controls and this association was also lost in FD patients. The genus *Exiguobacterium* has previously been identified in the duodenal microbiota (46) although its relationship with health is unclear. *Exiguobacterium* were strongly associated with the presence of Th2 effectors cells in the mucosa of FD patients but not controls. *Exiguobacterium* are highly adaptable microbes with some capable of degrading microplastics such as polystyrene (47). Given that microplastics have been suggested to contribute to DGBIs (48) and that microplastic exposure in humans can evoke a Th2 response (49, 50), it will be interesting to examine whether duodenal *Exiguobacterium* respond to microplastics in the diet.

Overall, the MAM of the control subjects without gut symptoms showed stronger associations with both peripheral and mucosal immune populations than the MAM of FD patients, supporting the concept of dysregulated mucosal homeostasis in FD. This included loss of associations between immune populations and *Veillonella* and *Prevotella* in FD patients, both of which have shown to have decreased relative abundances in FD patients (16). A notable exception to this overall trend was the genus *Cellulomonas*, which was positively correlated with Th2 populations and negatively correlated with Th17 populations in FD patients alone. Importantly, these T helper cell subsets have been identified as enriched in the duodenal mucosa of FD patients (3). *Cellulomonas* species have high capacity to catabolise antigenic gluten (51) and the balance of Th2/Th17 responses in FD patients have previously been linked to altered responses gluten stimulation (52). Increases in peripheral gut-homing T cells are also a common immunological feature in patients with DGBI (53). It is however unclear whether recruitment of these cells is driven by specific pathogens, food antigens or some other trigger. Differential digestion of potentially antigenic peptides from foods by microbes such as *Cellulomonas* may be a potential cause of these responses.

We attempted to understand whether the altered FD microbiota was associated with functional changes that may impact immune homeostasis in the gastrointestinal tract. Overall, the greatest changes were seen in the stomach, which may be driven by delayed gastric emptying commonly reported in FD (53, 54). Longer food residence times would potentially lead to greater metabolic breakdown of food, enriching related pathways in the stomach MAM. Indeed, we observed a number of pathways significantly increased within FD patients, such as nitrifier denitrification, mixed acid fermentation, and starch degradation V, associated with fermentation and carbohydrate degradation. Gas production is a significant component of fermentation end-products and we did not observe any pathways of gas utilisation (i.e., methanogenesis) that were significantly increased in FD patients, which may suggest that the excess gas production contributes to bloating, a common symptom reported in FD (25, 55). Within the duodenum, specific pathways associated with digestion, metabolic adaption, host colonisation and immune evasion were enriched in the FD patient MAM. Among the strongest associations were chondroitin sulfate degradation I which may be utilised by bacteria to reduce epithelial barrier function (43). Chondroitin sulfate is an extracellular matrix component that contributes to tight junction integrity (56), so increases in this pathway could impair normal gut barrier function. Other pathway associated with FD include protein N-glycosylation and CMP-pseudaminate biosynthesis which are related to microbial colonisation (57) and immune evasion (58), respectively. Collectively these pathways suggest that the microbiome’s capacity to disrupt normal homeostasis is altered in the FD MAM.

There are some limitations to this study, primarily the use of 16S rRNA amplicon sequencing of the MAM. This microbiome evaluation approach is limiting as any primer selection introduces bias and it does not provide strain resolution or functional capacity of the microbiome as shotgun metagenomics would. However, analysing the MAM by metagenomics is complicated by the overwhelming human DNA contamination of biopsy samples (59). Promisingly, recent research suggests gastrointestinal site specific culturomic based approaches are able to increase microbial yield of human biopsies suitable for shotgun metagenomic approaches [48, 49]. Unfortunately, these MAM samples were sequenced before this approach was available, there is no current culture medium for oesophageal MAM, and our biopsies were not collected in a cryopreservative solution appropriate for culturomics. While the number of samples in this study are modest, they are powered to identify the immune abnormalities in FD and our cohort exhibited the peripheral and mucosal immune alterations previously identified in FD patients (3, 11, 12).

In conclusion we have provided new evidence to support the concept that FD patients experience microinflammation and have an altered MAM of the upper gastrointestinal tract. Furthermore, to our knowledge this is the first study to investigate the FD immune-MAM associations and demonstrate a dysregulation of this relationship when compared to asymptomatic outpatient controls. Overall, our study supports the concept of a loss of host immune-microbiome homeostasis contributing to the pathology of FD. Changes in the MAM may favour the outgrowth of pathobionts that contribute to loss of “tolerance” to the normal microbiome constituents of the duodenum or lead to inappropriate digestion of antigen containing foods.

## Acknowledgements

The work was funded by a National Health and Medical Research Council Ideas grant and the Centre of Research Excellence in Digestive Health from the NHMRC. ECH is supported by the Australian NSW Health Round 5 Early-Mid Career Grant.

## AUTHOR CONTRIBUTIONS

ECH performed data collection, microbiome analysis, data interpretation and drafted the manuscript; GLB recruited patients, collected samples, and analysed histological and immune data; JB recruited patients and collected samples; SKang extracted samples for sequencing and with MM advised on analysis approaches and manuscript editing; SKeely planned and supervised research, reviewed data analysis and drafted manuscript; and NJT planned and supervised research, reviewed data analysis and manuscript. All authors have reviewed approved the final manuscript.

